# Identification of a small molecule that stimulates human β-cell proliferation and insulin secretion, and protects against cytotoxic stress in rat insulinoma cells

**DOI:** 10.1101/804807

**Authors:** Hans E. Hohmeier, Lu Zhang, Brandon Taylor, Samuel Stephens, Peter McNamara, Bryan Laffitte, Christopher B. Newgard

## Abstract

A key event in the development of both major forms of diabetes is the loss of functional pancreatic islet β-cell mass. Strategies aimed at enhancing β-cell regeneration have long been pursued, but methods for reliably inducing human β-cell proliferation with full retention of key functions such as glucose-stimulated insulin secretion (GSIS) are still very limited. We have previously reported that overexpression of the homeobox transcription factor Nkx6.1 stimulates β-cell proliferation, while also enhancing GSIS and providing protection against β-cell cytotoxicity through induction of the VGF prohormone. We developed an Nkx6.1 pathway screen by stably transfecting 832/13 rat insulinoma cells with a VGF promoter-luciferase reporter construct, using the resultant cell line to screen a 630,000 compound chemical library. We isolated three compounds with consistent effects to stimulate human islet cell proliferation. Further studies of the most potent of these compounds, GNF-9228, revealed that it selectively activates human β-cell relative to α-cell proliferation and has no effect on δ-cell replication. In addition, pre-treatment, but not short term exposure of human islets to GNF-9228 enhances GSIS. GNF-9228 also protects 832/13 insulinoma cells against ER stress- and inflammatory cytokine-induced cytotoxicity. In contrast to recently emergent Dyrk1a inhibitors that stimulate human islet cell proliferation, GNF-9228 does not activate NFAT translocation. These studies have led to identification of a small molecule with pleiotropic positive effects on islet biology, including stimulation of human β-cell proliferation and insulin secretion, and protection against multiple agents of cytotoxic stress.

## Introduction

Both major forms of diabetes involve loss of functional β-cell mass, but methods for reliably inducing human β-cell proliferation with full retention of key functions such as glucose-stimulated insulin secretion (GSIS) are still very limited. Early approaches focused on manipulation of oncogenes or core elements of the cell replication machinery such as cyclin D1, cdk6, p16 or p57 [1–3], but such strategies either led to functional impairment or were not therapeutically tractable due to the common use of these factors for cell cycle control in all tissues in the body. More recently, several groups reported that inhibition of the tyrosine kinase Dyrk1A with harmine or other small molecules stimulates human β-cell replication via activation of the NFAT/calcineurin pathway [4–6]. However, the wide tissue distribution of Dyrk1a suggests that inhibitors could trigger promiscuous cellular proliferation, and indeed, systemic administration of harmine activates islet α-, δ-, and ductal cell proliferation in addition to its effects on β-cells [5]. Therefore, additional strategies for expansion of functional human β-cell mass are needed, including mechanisms that are orthogonal to the Dyrk1A/NFAT pathway.

Our laboratory has performed a series of studies on overexpression of the homeobox transcription factor Nkx6.1 in pancreatic islets [7–12]. Key findings include: 1) Overexpression of Nkx6.1 in rat islets results in increased β-cell replication accompanied by improved GSIS [7, 8]; 2) Nkx6.1 induces expression of a prohormone, VGF, in both rat and human islets, which is processed to yield multiple peptides, including TLQP-21 [9]. TLQP-21 protects β-cells from apoptotic cell death, and VGF plays an important role in insulin vesicle trafficking [9, 12]. However, VGF or its peptides do not activate β-cell proliferation [9]; 3) Nkx6.1 activates β-cell replication by a pathway that is additive to and distinct from the pathway activated by overexpression of Pdx-1 [10, 11]. Importantly, overexpression of Nkx6.1 under control of the constitutive CMV promoter (causing expression in all islet cells) selectively stimulates proliferation of β-cells [8, 11], whereas overexpression of Pdx-1 via the same vector stimulates proliferation of both α- and β-cells via a secreted factor or factors [11, 13]. These results support the idea that Nkx6.1 activates a replication pathway with intrinsic β-cell specificity.

Based on these findings, we stably transfected the INS-1-derived 832/13 rat insulinoma cell line [14] with a plasmid containing the VGF promoter driving expression of luciferase, and used the resultant cell line to screen a 630,000 compound small molecule library. Given the positive impact of Nkx6.1 and VGF in islet cells [7–12], the screen was designed with a VGF-luc construct to capture small molecules that activate Nkx6.1 expression, stabilize or activate Nkx6.1 function, or serve as direct inducers of VGF. We ultimately isolated three compounds with robust and consistent effects to stimulate human islet cell proliferation. Further studies of the most potent of these compounds, GNF-9228, revealed that it selectively activates human β-cell relative to α- or δ-cell proliferation. Moreover, pre-treatment of human islets with GNF-9228 enhances insulin secretion. GNF-9228 also protects a rat β-cell line, INS-1 832/13, against cytotoxicity. Finally, GNF-9228 does not activate NFAT translocation, suggesting a mechanism of action distinct from Dyrk1a inhibitors.

## Materials and methods

### Cell culture and reagents

INS-1-derived 832/13 rat insulinoma cells were cultured as previously described [14]. Human islets were obtained from the Integrated Islet Distribution Program (IIDP) or from the University of Alberta Human Islet Core. Viability of human islets was tested by staining with propidium iodide as described in the IIDP SOP (http://iidp.coh.org), and were used only if viability was >75%. Human islets were cultured in RPMI-1640 (Invitrogen 11879) supplemented with 5.5 mmol/L glucose, 10% FCS (Sigma), 100 U/ml penicillin, 100 µg/ml streptomycin and 250 ng/ml amphotericin B (Gibco #15240). An adenovirus vector containing the cDNA for NFATc1 was obtained from Welgen, and stock solutions were diluted 1:8000 when added to Ins1 cells in 384 well plates.

### Generation of the 832/13-VGF-Luc Cell Line

Primers GAA GAT CTA TGG TCG AGGGCT GGC G and GGG GTA CCT GAC CCC CCT TCT CAG C corresponding to +77 to -2022 of the rat *VGF* promoter were used to amplify rat genomic DNA. The amplified PCR product was digested with BglII and KpnI for ligation upstream of a luciferase reporter gene that is contained in vector pGL4.21 (Promega). 832/13 cells were transfected with the resultant VGF promoter-luciferase reporter plasmid by electroporation and selected with puromycin [14] to yield stably transfected 832/13-VGF-Luc cells.

### Gene expression analyses

RT-PCR was used to measure the following transcripts, using the indicated Taqman probes: Rat Nkx6.1 : Rn01450076_m1, human NKX6.1: Hs00232355_m1, human VGF: Hs00705044_s1, human MYC: Hs00153408_m1. Gene expression was normalized against expression of cyclophilin A: human PPIA: Hs04194521_s1, rat Ppia: Rn00690933_m1.

### Use of 832/13-VGF-Luc cell line for small molecule high-throughput screen

High throughput screening was performed using 832/13-VGF-Luc cells. Briefly, 2,000 cells per well were plated in 5 µL RPMI in 1536-well plates. Individual molecules from a 630,000 compound chemical library assembled at the Genomics Institute of the Novartis Foundation (GNF) were added to single wells at a final concentration of 10 µM and incubated for 6 h. 2 µL of Bright Glo (Promega) was added to each well and luciferase-generated luminescence was measured with a plate reader (GNF Systems). Data were normalized to assay plate median and hits were called based on a robust deviation of 3. Hits in the primary screen were counterscreened against Ins1 cells stably transfected with a cyclic AMP response element (CRE)-luciferase reporter or HEK293T cells containing a PLTP-DR4B luciferase construct (pGL4.24 luciferase vector) [15]. The three compounds with human islet proliferative activity, GNF-9228, GNF-4088, GNF-1346 and the Dyrk1a inhibitor GNF-4877 [4] were dissolved in DMSO.

### Islet cell proliferation measured by EdU incorporation

For EdU labeling, a 1:1,000 dilution of EdU-labeling reagent (Invitrogen) was added to islet culture medium during the last 18h (Figure 3) or 72h (Figure 7) of cell culture. Islets were harvested and processed for immunofluorescence analyses as described previously [13]. Images were captured and analyzed with the Cellomics CX5 High Content (HC) cell based imaging system (Thermo) as described [13].

### Glucose-Stimulated Insulin Secretion

Human islets were pretreated with 10 µM GNF-9228 or DMSO for 72h. Insulin secretion was measured by static incubation of 30 islets in 4 replicate groups as previously described [9]. Secreted insulin was measured by ELISA (80-INSMR-CH10, ALPCO) and normalized to insulin content [9]. To measure the acute effects of GNF-9228, some batches of human islets were treated with GNF-9228 during the secretion assay only (no pre-treatment).

### Viability and cell toxicity assays

832/13 cells were seeded in 96-well tissue culture dishes. To induce cell toxicity, cells were treated with 200-500 nM thapsigargin or with a mixture of 1 ng/ml rat IL-1β (Sigma) + 100 U/ml rat IFN-γ (Thermo). Cell viability was measured using the CellTiter96 assay (Promega), or for apoptosis by measurement of caspase-3 cleavage [16].

### Pharmacokinetics of GNF-9228 in mice

The following studies were approved by the Duke University Institutional Animal Care and Use Committee (IACUC). We delivered 30 mg/kg of GNF-9228, GNF-1346, or GNF-4088 suspended in DMSO by intraperitoneal injection into normal C57BL6 mice and measured levels of the compounds in the blood by mass spectrometry over the following 24 hours. Six C57BL6 mice also received daily injections of 30 mg/kg GNF-9228 for one week, while 6 control mice received injections of DMSO. For all mice, BrdU was added to the drinking water at a concentration of 0.8 mg/ml. 24 h after the final GNF-9228 or DMSO injection, the mice were euthanized and pancreata were dissected, fixed in neutral-buffered formalin, and paraffin embedded. Slides were incubated overnight with guinea pig anti-insulin (Dako) and mouse anti-BrdU (Dako) antibodies, followed by detection with an AlexaFluor 488 conjugated goat anti-guinea pig and AlexaFluor 555 conjugated goat anti-mouse secondary antibody (Invitrogen), and counterstaining with DAPI. Images were captured and analyzed using OpenLab software. A minimum of 4 slides per pancreas spaced 75-100 μm apart were analyzed, comprising a total of approximately 10,000 cells per condition.

### Statistical Analyses

Data are presented as means + standard errors of mean (SEM) and graphed using GraphPad Prism 8.1 software. The differences between groups were compared using a Student’s t-test and values of P < 0.05 were considered significant.

## Results

### Cell-based screen for inducers of Nkx6.1 and VGF

We engineered a rat insulinoma (INS-1)-derived cell line, 832/13 [14], by stable transfection with a rat VGF promoter-luc reporter to generate a new cell line, 832/13-VGF-luc. 832/13-VGF-luc cells were screened with the 630,000 compound small molecule library assembled at the Genomics Institute of the Novartis Research Foundation (GNF). A total of 4437 compounds caused significant (standard deviation of three from the mean) stimulation of luciferase expression in the screen. These hits were counterscreened in HEK-293 cells stably transfected with an unrelated promoter-luciferase construct (PLTP-luc), or Ins1 cells transfected with a CREB-luciferase construct. The counterscreening protocol yielded 41 compounds with activity profiles similar to those shown for compounds GNF-9228 and GNF-7169, featuring robust, dose-dependent activation of the VGF-luc construct, with no activation of PLTP-luc (**Fig 1**). Since the VGF promoter contains a consensus cyclic AMP response element-binding protein (CREB) binding sequence [17], compounds were also counterscreened against a CREB-luc reporter. Compound GNF-7169 had no activity against the CREB-luc construct, whereas GNF-9228 had a weak activating effect (**Fig 1**).

**Fig 1.**
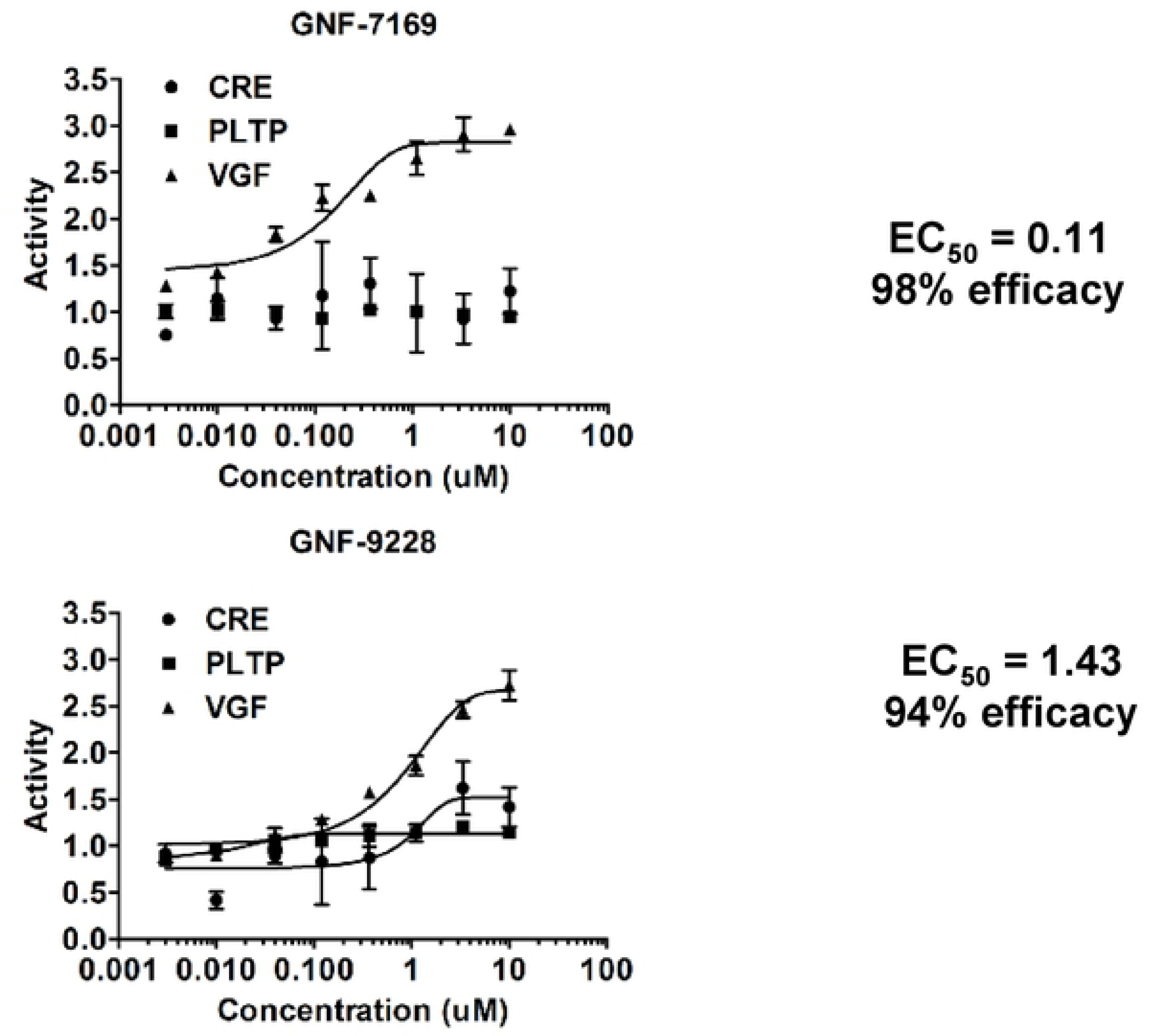
Representative “Hits” from small molecule screen. A high throughput screen was performed with 832/13 cells stably expressing a VGF promoter-luciferase reporter construct (832/13-VGF-luc cells). Hits were called based on a robust deviation of 3 from the mean. Hits were then confirmed in 8 point dose response studies in 832/13-VGF-Luc cells (VGF) and counter-screened against HEK-293T PLTP-Luc (PLTP) and Ins1 CRE-Luc (CRE) cell lines. Dose response curves for two representative hits, GNF-7169 and GNF-9228, are shown.

We studied the effects of the 41 lead compounds on islet cell proliferation, GSIS, and protection against apoptotic cell stress. Through this, we isolated three compounds (GNF-9228, GNF-1346, and GNF-4088) with robust and consistent effects to stimulate human islet cell proliferation. GNF-9228 and GNF-4088 are variants of a common chemotype, whereas GNF-1346 has a distinct structure (**Fig 2A**). Consistent with the design of the screen, both GNF-9228 and GNF-1346 caused potent induction of Nkx6.1 mRNA expression in 832/13 cells (**Fig 2B**). However, when tested on human islets, the two compounds caused a non-significant trend to increase Nkx6.1 mRNA, and had no effect on expression of VGF or the c-myc oncogene, a known activator of islet cell replication (**Figs 2C, 2D, 2E**) [18]. GNF-9228 was found to be the most potent of the three compounds for stimulation of human islet cell proliferation, and studies that follow therefore focus on the activities of this molecule.

**Fig 2.**
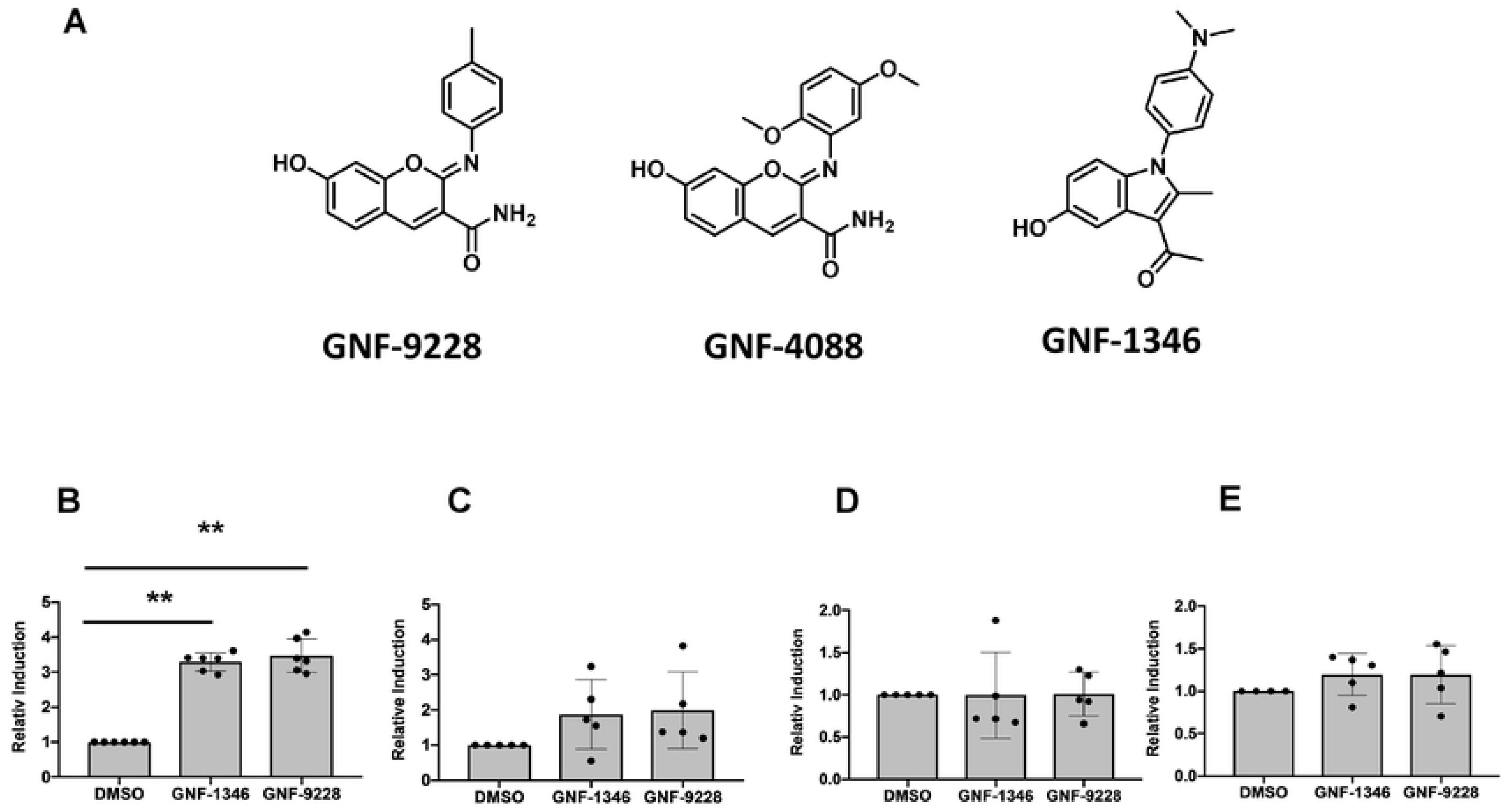
Structures of compounds GNF-9228, GNF-4088 and GNF-1346 and their effects on possible target genes. A. Structures of three compounds with human islet cell proliferative activity isolated from our 832/13-VGF-luc screen are shown. GNF-9228 and GNF-4088 have very similar structures, whereas GNF-1346 is distinct. **B. RT-PCR** measurement of Nkx6.1 levels in 832/13 cells following exposure to 10 µM GNF-1346, 10 µM GNF-9228 or vehicle (DMSO) for 6 hours. Five human islet samples from different donors were treated with 10 μM GNF-9228, 10 μM GNF-1346 or DMSO for 6 hours, followed by measurement of the following mRNAs by RT-PCR: **C.** Nkx6.1; **D.** VGF; **E.** c-myc. Data in panels B-E are expressed as fold-increase in 832/13 cells or islets treated with GNF-1346 or GNF-9228 relative to DMSO-treated, with n = 6 for 832/13 cells and n = 5 for human islet experiments (** p< 0.0001 comparing GNF-9228 or GNF-1346-treated cells to DMSO-treated cells in panel B).

### GNF-9228 selectively stimulates proliferation of human islet β-cells

Human islet cell replication was measured by EdU incorporation, and specific islet cell types were identified by insulin, glucagon, or somatostatin co-staining. A total of 7 human islet aliquots, each from a different donor, were tested for the effect of 10 µM GNF9228 on islet β-cell and α-cell replication. The dose of GNF-9228 was chosen as the maximally active dose based on titration studies in rat islets (**S1 Fig**). The individual human islet preps had substantial variation in the percent of β- and α-cells that were positive for EdU staining under control (DMSO-treated) conditions, including zero values for two of the α-cell experiments. The data are therefore presented as fold-response to normalize the differences in basal values. All 7 human islet preps exhibited an increase in β-cell and total cell EdU incorporation in response to GNF-9228 relative to DMSO treatment, with statistically significant average responses of 6-fold in total islet cells, and 7.3-fold in β-cells (**Fig 3**). The 5 human islet samples with detectable α-cell EdU staining at baseline all increased EdU incorporation into α-cells in response to GNF-9228, but here the average was 4.3-fold, and of marginal statistical significance (p=0.06). Additional information about the number of cells assayed and the percentages of EdU-positive islet cells, β-cells, and α-cells, is provided in (**S1 Table).** In addition, in a subset of human islet preparations with detectable co-staining of EdU and somatostatin under basal conditions, GNF-9228 caused no increase in EdU incorporation into δ-cells (**S2 Table**). Taken together, these data suggest that under the conditions used in these experiments (EdU exposure of 18 hours) GNF-9228 activates EdU incorporation into human islet cells in a β-cell selective fashion, with a marginal effect on α-cells and no detectable effect on δ-cells.

**Fig 3.**
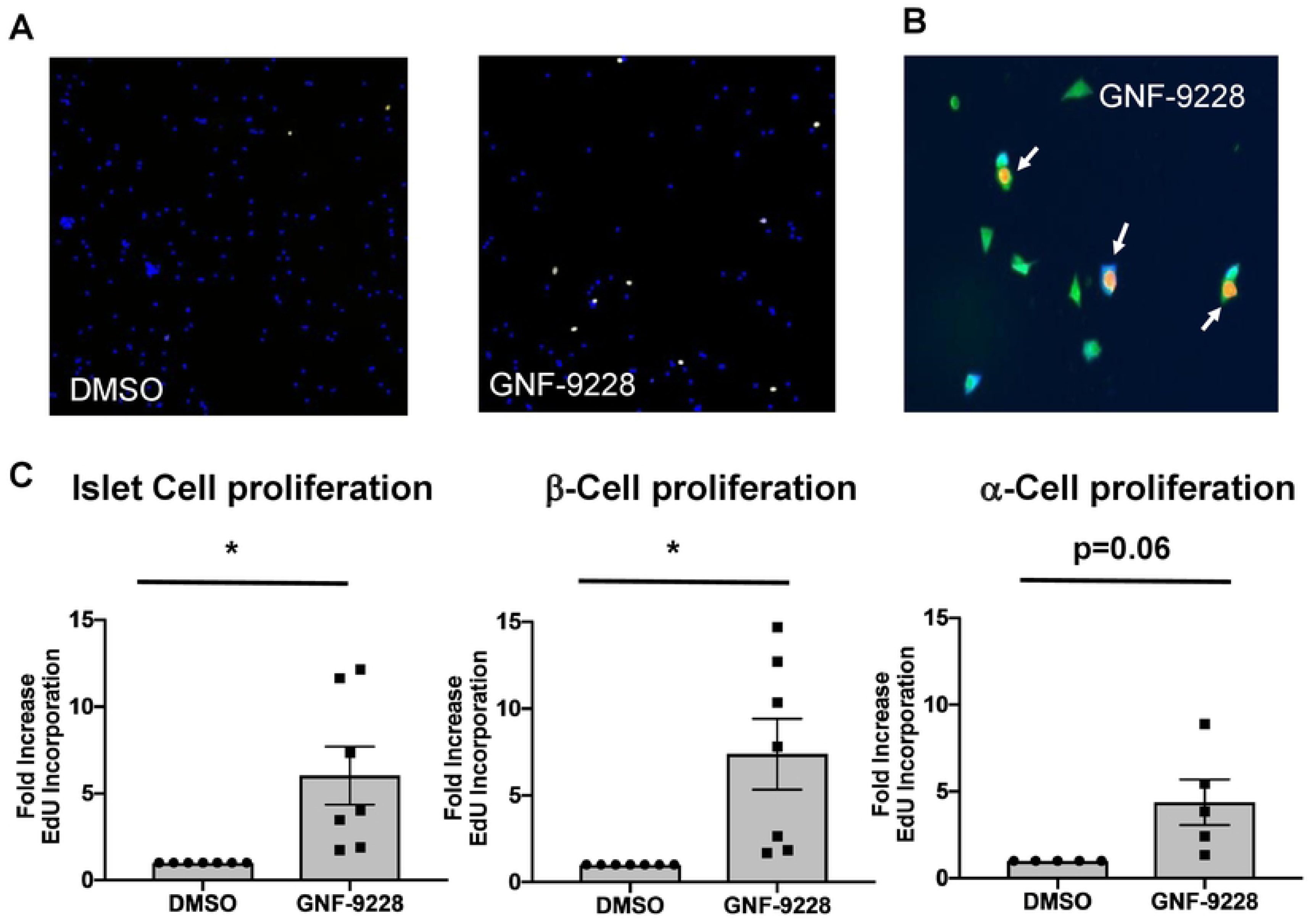
GNF-9228 selectively stimulates human Islet β-cell replication. Intact human islets were cultured for 72 h in the presence of 10 µM GNF-9228 or DMSO (vehicle control). EdU was added for the last 18 h of culture. Islets were dispersed, stained for EdU incorporation and treated with antibodies against EdU, insulin or glucagon, and immunofluorescent signals were detected and quantified with a Thermo Scientific Cellomics CX5-High Content (HC) cell imaging system**. A.** Low magnification image showing increase in islet cell EdU incorporation (yellow) in GNF-9228 versus DMSO-treated cells, with cells visualized by DAPI staining (blue). **B.** High magnification image showing two Edu (orange nuclei), insulin (green cytoplasm) co-positive cells and one Edu (orange nucleus), glucagon (blue cytoplasm) co-positive cell. **C.** Bar graph summary of data expressed as the fold-increase in EdU positive cells across all islet cells assayed (left), in β-cells (EdU and insulin co-positive cells (center), and α-cells (EdU and glucagon positive cells (right), representing experiments performed on 7 independent human islet aliquots (total cells and β-cells) and 5 human islet samples (α-cells), expressed as mean +/- S.E.M.(* p < 0.05). See S1 Table for further details.

### GNF-9228 stimulates insulin secretion in rat and human islets

We next tested the impact of GNF-9228 on insulin secretion. Acute (less than 24 hour exposure) of islets to GNF-9228 had no impact on GSIS (**S1 Fig**). However, pre-exposure of human islets to 5 or 10 µM GNF-9228 for 72 hours caused a 75-140% increase in insulin secretion in the presence of stimulatory (16.7 mM) glucose compared to human islets treated with DMSO (**Fig 4**). Insulin secretion was normalized to insulin content in these experiments, which showed a non-significant trend to decrease in response to GNF-9228 treatment (human islets treated for 72 h with 5 µM or 10 µM GNF-9228 had 86 ± 9.1% and 83 ± 12.9% of the insulin content measured in DMSO-treated islets, respectively). This non-significant change in insulin content does not explain the enhanced insulin secretion elicited by GNF-9228 treatment. We also note that insulin secretion at basal (2.8 mM) glucose levels increased significantly at 5 µM, and showed a non-significant trend to increase at 10 µM GNF-9228 (**Fig 4**). This increase in basal insulin secretion caused the stimulation index (insulin secreted at 16.7 mM relative to 2.5 mM glucose) to be essentially identical in the GNF-9228 and DMSO-treated islets. Nevertheless, pretreatment with, but not acute exposure to GNF-9228 enhances insulin secretion at stimulatory glucose in human islets.

**Fig 4.**
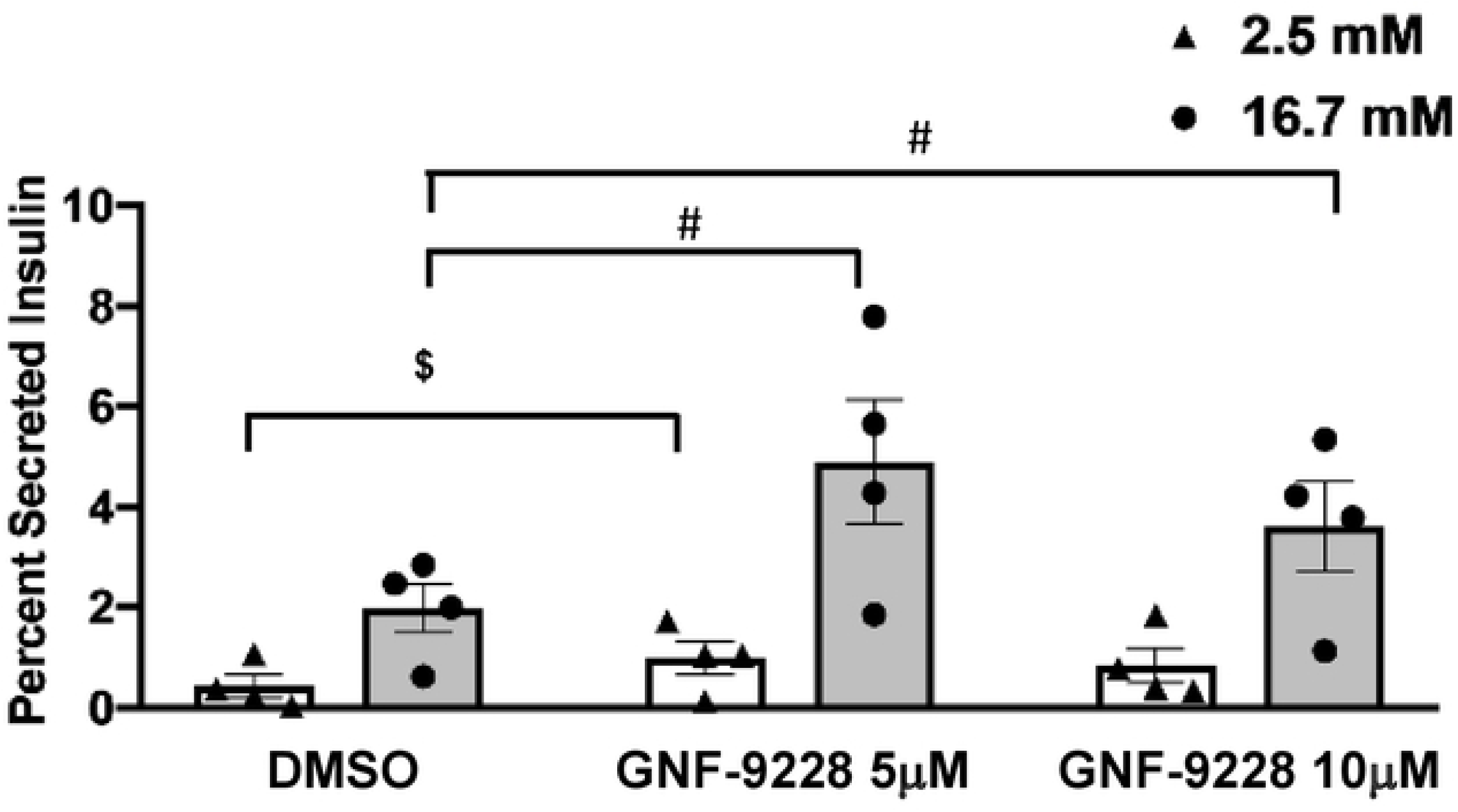
Pre-treatment of human islets with GNF-9228 enhances insulin secretion. Intact human islets were pre-treated for 72 h with 5 or 10 µM GNF-9228 or DMSO (vehicle control). Glucose-stimulated insulin secretion was then measured by static incubation in secretion buffer containing 2.5 mM glucose followed by 16.7 mM glucose for 1 h each. Secreted insulin is reported as a percentage of insulin content. Data represent the mean +/- S.E.M. from 4 independent human islet preparations, each assayed in quadruplicate. (# p < 0.005 compared to DMSO treated cells at 16.7 mM glucose; ^$^ p < 0.01 compared to DMSO treated cells at 2.5 mM glucose).

### GNF-9228 protects 832/13 cells from ER stress and cytokine-induced cytotoxicity

To test GNF-9228 for its potential pro-survival effects, 832/13 cells were exposed to 500 nM thapsigargin (TG) for 6 hours in the presence and absence of 10 µM GNF-9228. Co-treatment of 832/13 cells with TG + GNF-9228 resulted in a clear reduction in cleaved caspase-3 compared to control cells incubated with TG + DMSO (**Figs 5A and 5B**). In another set of experiments, 832/13 cells were pretreated with GNF-9228 or vehicle for 24 h followed by 24 h exposure to 200 or 250 nM thapsigargin (TG) or a mixture of the cytotoxic cytokines IL-1β and γ-IFN in the presence or absence of GNF-9228 (**Fig 5C**). In these experiments, cell viability was measured with the mitochondrial activity dye MTS. Exposure to 250 nM TG + DMSO decreased cell viability by 67%, whereas cells treated with TG + GNF-9228 suffered only a 31% reduction in viability. Similarly, exposure of vehicle-treated 832/13 cells to the inflammatory cytokines IL-1β + γ-IFN caused a 67% reduction in viability, whereas GNF-9228 limited the effect to a 40% reduction. Thus, GNF-9228 exhibits anti-apototic and pro-cell survival effects in the INS-1-derived 832/13 cell line. Note that we chose to perform our experiments in these cells based on our finding that they are far more sensitive to activation of apoptosis than primary islet cells [16], suggesting that the cell line provides the most stringent model for demonstrating cytoprotective effects of GNF-9228 at this stage of the work.

**Figure 5.**
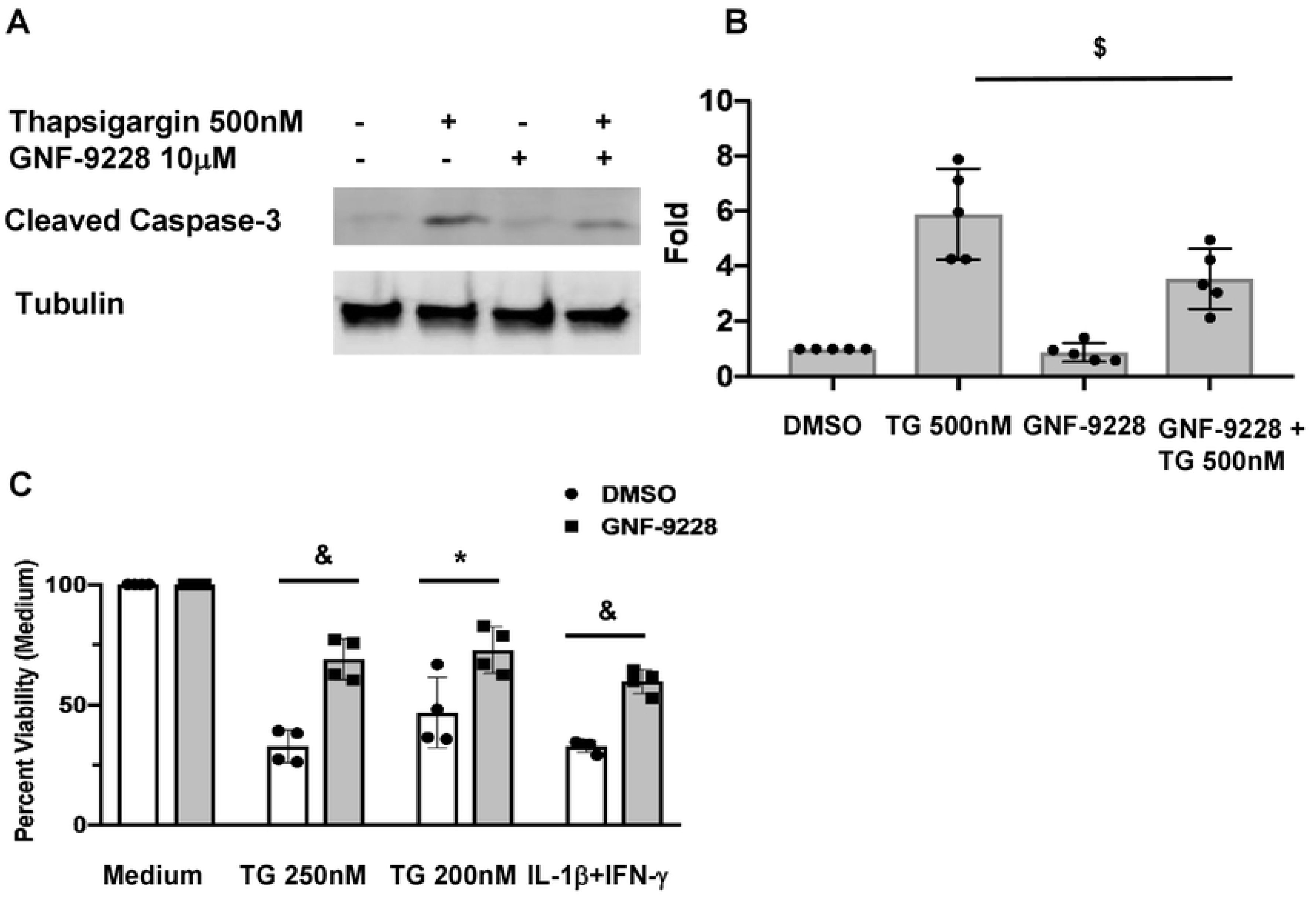
GNF-9228 protects 832/13 cells from ER stress and cytokine-induced cytotoxicity. A. 832/13 cells were treated with 500 nM thapsigargin (TG) for 6h in the presence and absence of 10 µM GNF-9228. A representative immunoblot of cleaved caspase-3 is shown. **B.** Bar graph summary of 4 independent densitometric measurements of cleaved caspase-3 in 832/13 cells exposed to 500 nM TG ± 10 µM GNF-9228 or DMSO for 6 hours. **C.** 832/13 cells were pre-treated with 10 µM GNF-9228 or DMSO for 24 h followed by 24 h incubation with two doses of thapsigargin (TG) or a mixture of IL-1β and γ-IFN. Cell viability was measured by MTS assay. (^&^p< 0.001; * p < 0.05; ^$^ p < 0.01, comparing GNF-9228 to DMSO-treated cells).

### Comparison of the effects of GNF-9228 and the Dyrk1A inhibitor GNF-4877

To test whether GNF-9228 activates islet cell proliferation by a mechanism similar to that reported for Dyrk1a inhibitors [4–6], we compared the effects of GNF-9228 and GNF-1346 to those of GNF-4877 [4] and harmine [5] on NFAT nuclear translocation. Treatment with increasing doses of these agents demonstrated clear activation of NFAT nuclear translocation by GNF-4877 and harmine, but not GNF-9228 or GNF-1346 (**Figs 6A and 6B**). Consistent with these findings, the robust proliferative effect of GNF-9228 in rat islets was not affected by co-treatment with cyclosporin A, a potent NFAT pathway inhibitor (**S3 Fig**). Also indicative of a unique mechanism of action, GNF-9228 caused a significant 3.9-fold increase in VGF-luc activity relative to DMSO treatment (p < 0.005), whereas GNF-4877 failed to cause significant luc activation in this assay (**Fig 6C**).

**Figure 6.**
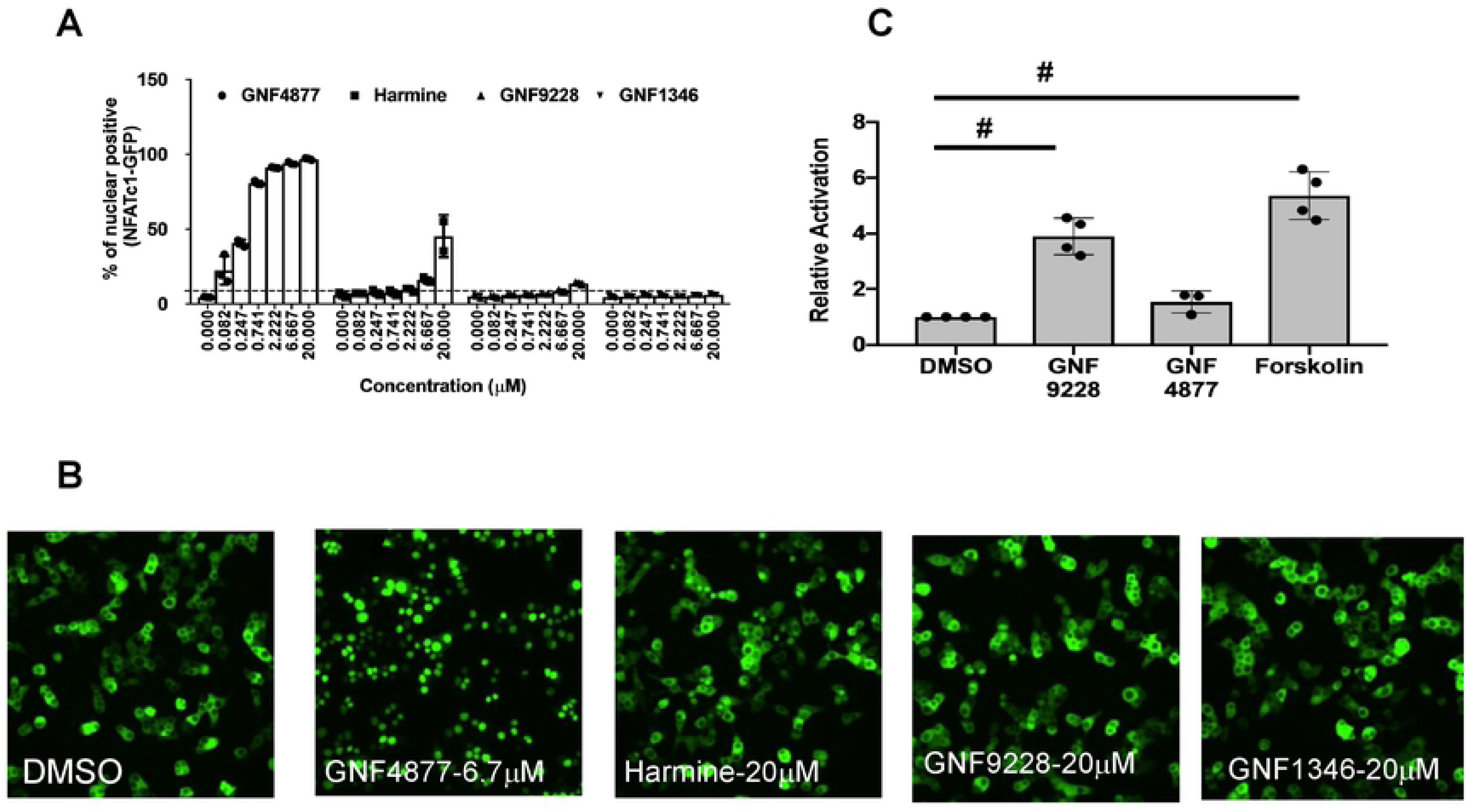
GNF-9228 and GNF-1346 signal by distinct mechanisms relative to Dyrk1a inhibitors. A. Ins1 cells treated with an NFATc1-GFP adenovirus were treated with a range of doses of the Dyrk1a inhibitors GNF-4877 or harmine, GNF-9228, or GNF-1346 or DMSO for 2h. Following treatment, cells were fixed and imaged for GFP nuclear localization. Data is quantified as percent of cells with nuclear localization of GFP for n = 3 experiments. **B.** Representative images showing subcellular localization of NFATc in the presence of 10 µM GNF-4877, harmine, GNF-9228, GNF-1346, or DMSO; **C.** 832/13 cells were transiently transfected with the VGF-luc plasmid used to generate 832/13-VGF-luc cells and treated with 10 µM GNF-9228, 5 µM GNF-4877, or 5 µM forskolin for 24 h followed by measurement of luc-generated luminescence. Data are expressed as fold-change relative to DMSO-treated cells in four independent experiments (^#^ p<0.005).

We also compared the proliferative effects of 10 µM GNF-9228 to those of 5 µM GNF-4877, a concentration reported as maximally active for islet cell proliferation [4]. In these experiments (**Figure 7**) human islets were exposed to EdU for 72 hours, in contrast to the studies in Figure 3 where EdU was provided for 18 hours. All 6 human islet preps responded to GNF-9228 or GNF-9228 + GNF-4877 with an increase in EdU incorporation into total islet cells and β-cells, with statistically significant average responses of approximately 10-fold to both treatment conditions (**Fig 7 and S3 Table**). GNF-4877 also increased EdU incorporation into β-cells in all 6 human islet preps, with an average increase of 4-fold (p = 0.06). (**Fig 7**). Combining the two compounds had no additive effect on total cell or β-cell proliferation. GNF-9228 or GNF-9228 + GNF-4877 also caused an average, statistically significant 5-fold increase in α-cell EdU incorporation in 5 separate human islet aliquots (**Fig 7 and S3 Table**). GNF-4877 alone also increased α-cell EdU incorporation in all 5 preps surveyed, with an average 3-fold increase (p = 0.06). We also measured EdU incorporation into δ-cells in human islet preps treated with EdU for 72 hours and again found no consistent effect of GNF-9228 (**S1 Table**). Thus, with longer duration of EdU exposure, we find that GNF-9228 increases both β-cell and α-cell but not δ-cell replication, with a preferential effect on β-cells.

**Fig 7.**
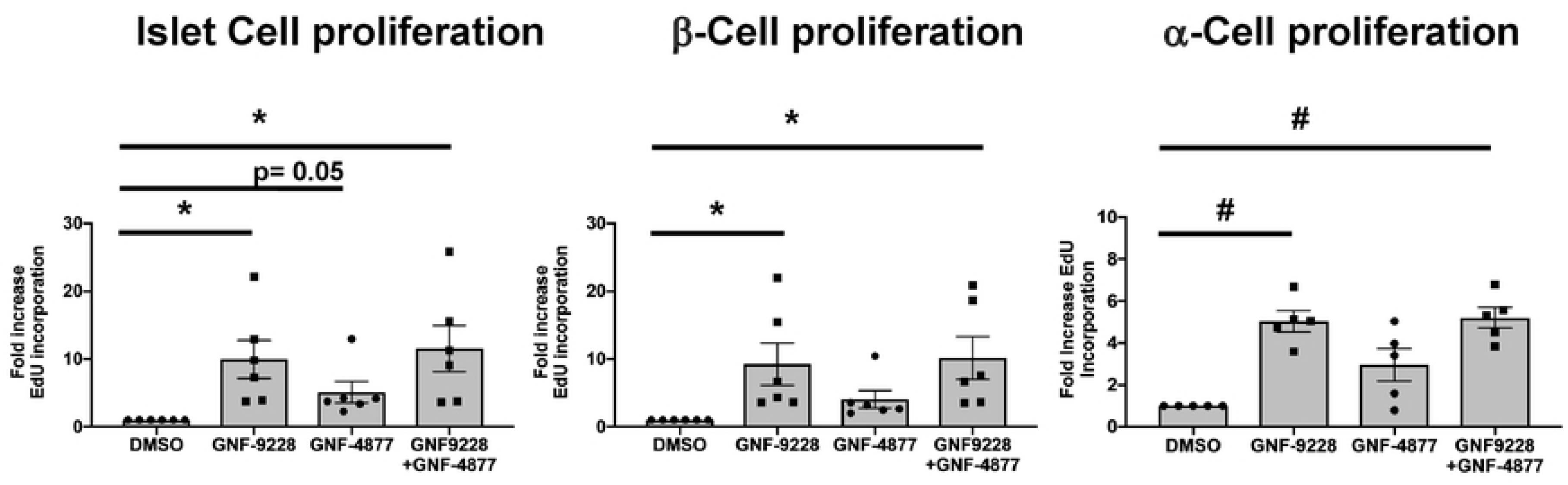
Comparison of effects of GNF-9228 and the Dyrk1a inhibitor GNF-4877 on human islet cell proliferation. Intact human islets were cultured for 72h in the presence of 10 µM GNF-9228, 5 µM GNF-4877, a combination of both, or DMSO (vehicle control). EdU was added for the last 72 h of culture. Islets were dispersed, stained for EdU incorporation and treated with antibodies against insulin or glucagon, and immunofluorescent signals assayed with a Thermo Scientific Cellomics CX5-High Content (HC) cell imaging system. Data are expressed as the fold-increase in total EdU positive cells (left), β-cells (EdU and insulin co-positive cells) (center), and α-cells (EdU and glucagon positive cells) (right), representing experiments performed on 6 independent human islet aliquots (total cells and β-cells) and 5 human islet samples (α-cells), expressed as mean +/- S.E.M. (* p< 0.05, ^#^ p < 0.005 relative to DMSO control). See S3 Table for more details.

### Compounds GNF-9228, GNF-1346, and GNF-4088 have poor *in vivo* pharmacodynamic properties

To test the effects of our compounds in the *in vivo* setting, we first injected 30 mg/kg of GNF-9228 suspended in DMSO into normal C57BL6 mice and measured its blood levels over the following 24 hours. Although high levels of GNF-9228 were detected at the earliest time points (Cmax = 8493 nM), the compound was cleared rapidly, with levels falling to 853 nM at 1 hour post-injection (half life = 8 minutes; **S4 Fig**). The pharmacokinetic properties of GNF-1346 and GNF-4088 were even less encouraging. Nevertheless, we performed a study in which GNF-9228 was injected into C57BL6 mice daily at a dose of 30 mg/kg for one week, with mice receiving BrdU in the drinking water at a concentration of 0.8 mg/ml throughout the study. GNF-9228 treatment caused no incease in BrdU incorporation into mouse islet cells compared to DMSO-treated mice, likely due to its rapid clearance.

## Discussion

Both major forms of diabetes involve loss of functional β-cell mass. In type 1 diabetes (T1D), islet β-cells are destroyed by an autoimmune mechanism, whereas in type 2 diabetes (T2D), the decline in β-cell mass and function is mediated by metabolic fuel overload and inflammatory pathways [19]. Thus, for both T1D and T2D, strategies for regeneration of functional β-cell mass should ideally address all of the contributory elements of β-cell failure, including susceptibility to cytotoxic agents, loss of proliferative capacity, and impairment of insulin secretion. In addition, the strategy chosen should target regeneration of islet β-cells specifically, or at least selectively, with minimal activity against other islet or peripheral cells. In light of these challenges, it is not surprising that no β-cell regenerative drugs currently exist for humans with diabetes.

The current study was built upon work showing that overexpression of the homeobox transcription factor Nkx6.1 stimulates islet β-cell replication in a selective fashion, while also enhancing GSIS [7–12]. VGF is a gene that is robustly induced by Nkx6.1 overexpression, and subsequent studies demonstrated several salutary effects of the VGF prohormone and its encoded peptides such as TLQP-21 on β-cell survival and function [9, 12]. Based on these findings, we designed a cell-based screening strategy involving stable transfection of 832/13 cells with a VGF promoter-luciferase reporter gene. The screen yielded three compounds representing two chemotypes (Fig 2) with a robust capacity to stimulate human islet cell replication.

The most potent of these compounds, GNF-9228, has a remarkable combination of properties. First, it selectively activates human β-cell EdU incorporation, with a lesser effect on α-cell replication and no effect on δ-cells. Second, pre-exposure of human islets to GNF-9228 enhances insulin secretion at stimulatory glucose levels (16.7 mM). Third, pre-treatment of 832/13 rat insulinoma cells with GNF-9228 is cytoprotective against the ER stress-inducing agent thapsigargin or a mixture of the cytotoxic cytokines IL-1β + γ-IFN. We recognize that the compound is not entirely β-cell specific, with a lesser, but significant effect to stimulate α- cell replication (Figure 7). We also acknowledge that the compound tends to increase insulin secretion at basal glucose levels, an effect that if mimicked in the *in vivo* setting could result in increased risk for hypoglycemia. Further evaluation of these potential shortcomings will require development of GNF-9228 analogs with improved *in vivo* pharmacodynamics.

In addition to our work on Nkx6.1 and Pdx-1 as upstream inducers of islet cell proliferation [7–13], other signaling pathways that have been identified with potential for activating human β- cell proliferation include the PDGF pathway [21], signaling by TGF-β family members [13,20,22], and glucose-regulated activation of proliferation mediated by ChREBP and c-myc [17, 23]. Pathways that regulate translocation of the NFAT transcription factor have also received attention stemming from early studies showing that conditional ablation of NFAT caused a reduction of β-cell mass in mouse models [24–26]. The tyrosine kinase Dyrk1A is a potent regulator of NFAT translocation [24]. Recently, Dyrk1A inhibitors with islet cell proliferative effects have emerged including harmine, identified in a small molecule screen for c-myc inducers [5], GNF-4788, synthesized as a derivative of an aminopyrazine scaffold previously shown to have proliferative activity [4, 27], and 5-IT, an adenosine kinase inhibitor that cross-reacts with Dyrk1a [6]. All of these agents cause substantial increases in human β-cell proliferation, with either no impairment of function [4, 5], or an actual increase in GSIS with prolonged (12 days) treatment [6].

While these results provide encouragement for further investigation of Dyrk1a inhibitors for diabetes therapy, concerns about their ultimate practical utility include: 1) When added to human islets, harmine caused a significant increase in proliferation of non-β-cells, including α-cells, δ-cells and ductal cells [5]. This coupled with the broad expression of Dyrk1a suggests that unwanted proliferative responses might occur in peripheral tissues in response to systemic administration of Dyrk1a inhibitors; 2) Harmine activates c-myc, explaining at least a portion of its replicative activity [5], GNF-4788 inhibits the Dyrk1a-related kinase GSK-β [4], and 5-IT was originally isolated as an adenosine kinase inhibitor [6], suggesting possible off-target effects of these agents; 3) Overexpression of c-myc has independent effects on human islet cell proliferation [17], but also appears to drive islet de-differentiation and impairment of GSIS [28]; 4) In contrast to our work with GNF-9228, none of the studies on Dyrk1a inhibitors report a cytoprotective effect.

Importantly, our studies suggest that GNF-9228 activates signaling pathways distinct from those used by Dyrk1a inhibitors. Unlike GNF-4788 and harmine, GNF-9228 does not induce nuclear translocation of an NFAT-GFP fusion gene. Moreover, the proliferative effects of GNF-9228 are not blocked by NFAT translocation inhibitors such as cyclosporin A, and GNF-9228 but not GNF-4788 activates the VGF-luc reporter in 832/13-VGF-luc cells. However, GNF-9228 and GNF-4788 have no additive effects on human islet cell proliferation, possibly suggesting some mechanistic commonality of these agents that remains to be defined.

Although we have identified differences in GNF-9228 and Dyrk1a inhibitor cell signaling, we have not yet defined the mechanism(s) of action of GNF-9228. In human islets, a dose of GNF-9228 (10 µM) that activates β-cell proliferation and GSIS causes a non-significant trend for increase of Nkx6.1 mRNA and has no effect on VGF or c-myc mRNA. This suggests that Nkx6.1 does not mediate the replicative effect of GNF-9228, and VGF is unlikely to explain its effects on insulin secretion and cell survival. Further studies will be required to discern the mechanism(s) of action of GNF-9228.

In sum, we report the identification of a small molecule that stimulates human β-cell proliferation, enhances insulin secretion, and protects against cytotoxic agents. An obvious next step is to test the impact of GNF-9228 on islet cell proliferation, function, and survival *in vivo*, but unfortunately all three of the compounds that emerged from our screen have poor pharmacodynamic properties. Our current focus is on development of chemically modified versions of GNF-9228 with enhanced bioavailability to allow *in vivo* testing. Further studies are also required to understand its mechanism(s) of action.

## Acknowledgements

These studies were supported by a grant from the Juvenile Diabetes Research Foundation, 1-SRA-2017-356. The authors thank Drs. Patricia Kilian and Andrew Rakeman for helpful discussion and suggestions, and Lisa Poppe and Danhong Lu for excellent technical assistance. We also thank the Integrated Islet Distribution Program and Dr. Patrick MacDonald and the University of Alberta Islet Core Laboratory for provision of human islets for our studies.

